# Genetic Single Neuron Anatomy reveals fine granularity of cortical interneuron subtypes

**DOI:** 10.1101/219485

**Authors:** Xiaojun Wang, Jason Tucciarone, Siqi Jiang, Fangfang Yin, Bor-shuen Wang, Dingkang Wang, Yao Jia, Xueyan Jia, Yuxin Li, Tao Yang, Zhengchao Xu, Masood A. Akram, Yusu Wang, Shaoqun Zeng, Giorgio A. Ascoli, Partha Mitra, Hui Gong, Qingming Luo, Z. Josh Huang

## Abstract

Parsing diverse nerve cells into biological types is necessary for understanding neural circuit organization. Morphology is an intuitive criterion for neuronal classification and a proxy of connectivity, but morphological diversity and variability often preclude resolving the granularity of discrete cell groups from population continuum. Combining genetic labeling with high-resolution, large volume light microscopy, we established a platform of genetic single neuron anatomy that resolves, registers and quantifies complete neuron morphologies in the mouse brain. We discovered that cortical axo-axonic cells (AACs), a cardinal GABAergic interneuron type that controls pyramidal neuron (PyN) spiking at axon initial segment, consist of multiple subtypes distinguished by laminar position, dendritic and axonal arborization patterns. Whereas the laminar arrangements of AAC dendrites reflect differential recruitment by input streams, the laminar distribution and local geometry of AAC axons enable differential innervation of PyN ensembles. Therefore, interneuron types likely consist of fine-grained subtypes with distinct input-output connectivity patterns.

## Introduction

Diverse nerve cells wire up intricate neural circuits that underlie animal behavior. Defining and cataloging the basic elements, groups of neurons that share anatomical, physiological and molecular properties are necessary for understanding the organizational logic of neural circuits (Huang and Zeng, 2013; Lerner et al., 2016). As phenotypic variations of neurons often span substantial parameter space and appear continuous as well as discrete, it is necessary to carry out comprehensive, quantitative, and scalable single cell analysis (instead of tissue level analysis of mixed cell populations) to resolve the appropriate granularity of cell type definition (Zeng and Sanes, 2017). Recent advances in single cell RNA sequencing (scRNAseq) enable comprehensive and quantitative measurements of cellular transcriptome profiles at massive scale, and computational analyses reveal increasing number of “transcriptional types” and discrete as well as continuous variations (Macosko et al., 2015; Paul et al., 2017; Poulin et al., 2016; Tasic et al., 2016; Zeisel et al., 2015). As neuronal phenotypes are inherently multi-modal, it is necessary to achieve single cell analyses across multiple cell features toward an integrated definition of neuron types that encapsulate the issue of granularity.

Neuronal morphology has been an intuitive first level description of cell features and types since Cajal’s initial observation. In several invertebrate systems (Aso et al., 2014; Chiang et al., 2011) and the vertebrate retina (Sanes and Masland, 2015) in which neurons are relatively small and stereotyped, comprehensive and quantitative single neuron morphometry has allowed operational and consensual definition of neuron types. Morphology based cell catalogues in these systems have been achieved or within reach (Aso et al., 2014; Hobert et al., 2016; Seung and Sumbul, 2014), which provide a foundation for multi-modal analysis and for exploring neural circuit organization. In the mammalian brain, however, the vast diversity, large spatial span (e.g. 100nm axon extending centimeters), and seemingly endless variations of neuronal shapes present unique challenges in morphological tracing and analysis (Huang and Zeng, 2013; Lichtman and Denk, 2011). Single neuron anatomy in the mammalian brain requires overcoming several technical hurdles. The first is labeling: to systematically, reliably, sparsely and completely label specific sets of individual neurons. The second is imaging: to achieve axon resolution imaging in brain-wide volume (Economo et al., 2016; Gong et al., 2013; Li et al., 2010). The third is cell reconstruction: to convert large image stacks into digital datasets of single neuron morphology. The fourth is analysis: to register neuronal morphology within appropriate spatial coordinate framework, and to extract, quantify and classify biologically relevant attributes (e.g. those relate to neural connectivity).

Here, we have established a robust genetic Single Neuron Anatomy (gSNA) platform in the mouse that overcomes some of these challenges. We designed a genetic method to achieve specific, sparse, complete, and reliable cell labeling. Combined with dual-color fluorescence Micro-Optic Sectioning Tomography (dfMOST; (Gong et al., 2016), this enabled axon resolution and brain-wide imaging and spatial registration of genetically targeted neurons. We focused our analysis on one of the most distinctive cortical GABAergic interneurons – the axo-axonic cells (AACs) that specifically innervate the axon initial segment (AIS) of glutamatergic pyramidal neurons (PyNs) (Somogyi et al., 1982; Taniguchi et al., 2013) and likely control spike initiation. Complete reconstruction of single AACs and their precise registration along cortical laminar coordinate allowed quantitative analysis of AAC morphology in the context of input-output connectivity. We discovered that cardinal AACs consists of multiple discrete subtypes distinguished by their laminar position, dendritic and axonal arborization pattern, and geometric features. The laminar arrangements of AAC dendrites may allow differential recruitment by presynaptic input streams. Furthermore, the laminar stratification of AAC axon arbors correlates with the distribution of PyN subsets and the local geometry of AAC axon terminals differentially conform to the laminar features of PyN AIS, suggesting AAC subtypes that differentially innervate PyN ensembles. Our results support a hierarchical scheme of neuronal classification (Zeng and Sanes, 2017) and suggest that cardinal neuron types consist of fine-grained subtypes, which can be deduced from light microscopy and mesoscale analyses that inform input-output connectivity patterns. The gSNA platform enables scalable and comprehensive single neuron anatomical analysis, which will provide foundational datasets for neuron type definition, classification, and census in the mammalian brain.

## Results

### Establishing a genetic single neuron anatomy (gSNA) platform

Our gSNA platform consists of four components (**Fig. 1a**). The first is a method to systematically label different sets of genetically targeted individual neurons to their entirety; the second is a technology for simultaneous imaging of labeled neurons at axon resolution and all other cell body positions throughout the entire mouse brain (dfMOST) (Gong et al., 2016); the third is a procedure to completely reconstruct single neurons from brain volume image stacks; the fourth is an analysis pipeline that registers and quantifies neuronal morphology within an appropriate spatial coordinate system that reflect network connectivity.

**Figure 1.**
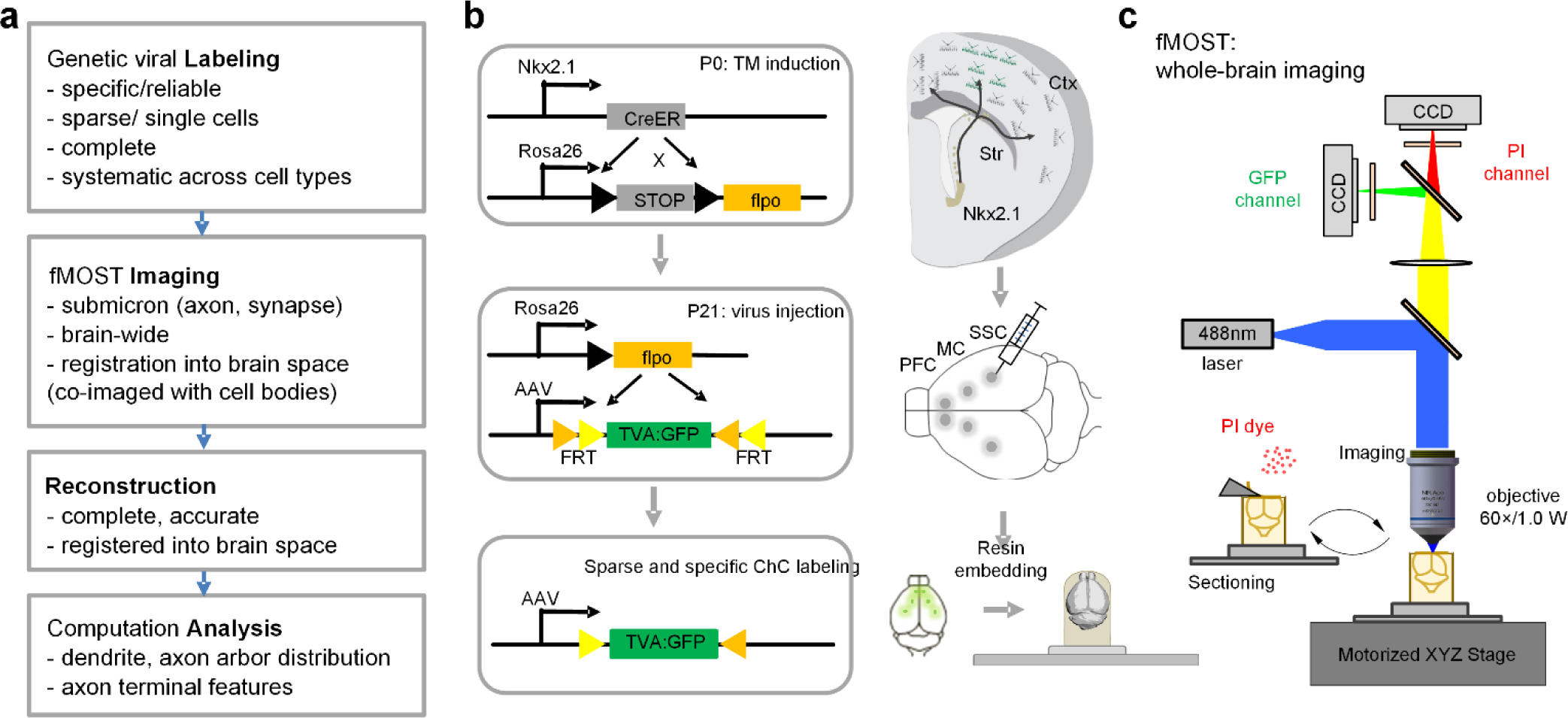
Schematic of the genetic Single Neuron Anatomy (gSNA) platform applied to the mouse brain. (**a**) Pipeline and components of gSNA. (**b**) Left: scheme of genetic and viral strategy for the labeling of axo-axonic cells (AACs). A transient CreER activity in MGE progenitors is converted to a constitutive Flp activity in mature AACs. Flp-dependent AAVs injected in specific cortical areas enables sparse and robust AAC labeling. (**c**) fMOST high resolution whole-brain imaging. Two-color imaging for the acquisition of GFP (green) channel and PI (propidium iodide, red) channel signals. PI stains brain cytoarchitecture in real time, therefore provides each dataset with a self-registered atlas. A 488 nm wavelength laser was used for the excitation of both GFP and PI signals. Whole-brain coronal image stacks were obtained by sectioning (with a diamond knife) and imaging cycles at 1 μm z-steps, guided by a motorized precision XYZ stage.

Cell labeling is the starting point of neuroanatomy, yet specific, sparse, complete, and systematic labeling of diverse nerve cells has been an enduring challenge since the invention of Golgi stain. We combined genetic and viral methods to solve this problem in the mouse brain in several steps. We first generated gene knockin recombinase driver lines that allow specific and reliable targeting of cell populations defined by single or combinatorial gene expression (He et al., 2016; Taniguchi et al., 2011). Next, inducible CreER drivers enable titration of sparseness for single cell labeling (He et al., 2016). Further incorporation of recombinase-activated AAV vectors achieves high level reporter gene expression for complete cell labeling (**Fig. 1b, Fig. 2**). Systematic iterations of this strategy with the accumulation of driver lines (e.g. similar to those in Drosophila; (Jenett et al., 2012)) will enable increasingly comprehensive coverage of genetically defined populations across the mouse brain. Here we demonstrate the gSNA platform in resolving morphological diversity and granularity by analyzing a well-recognized interneuron type in the cerebral cortex.

**Figure 2.**
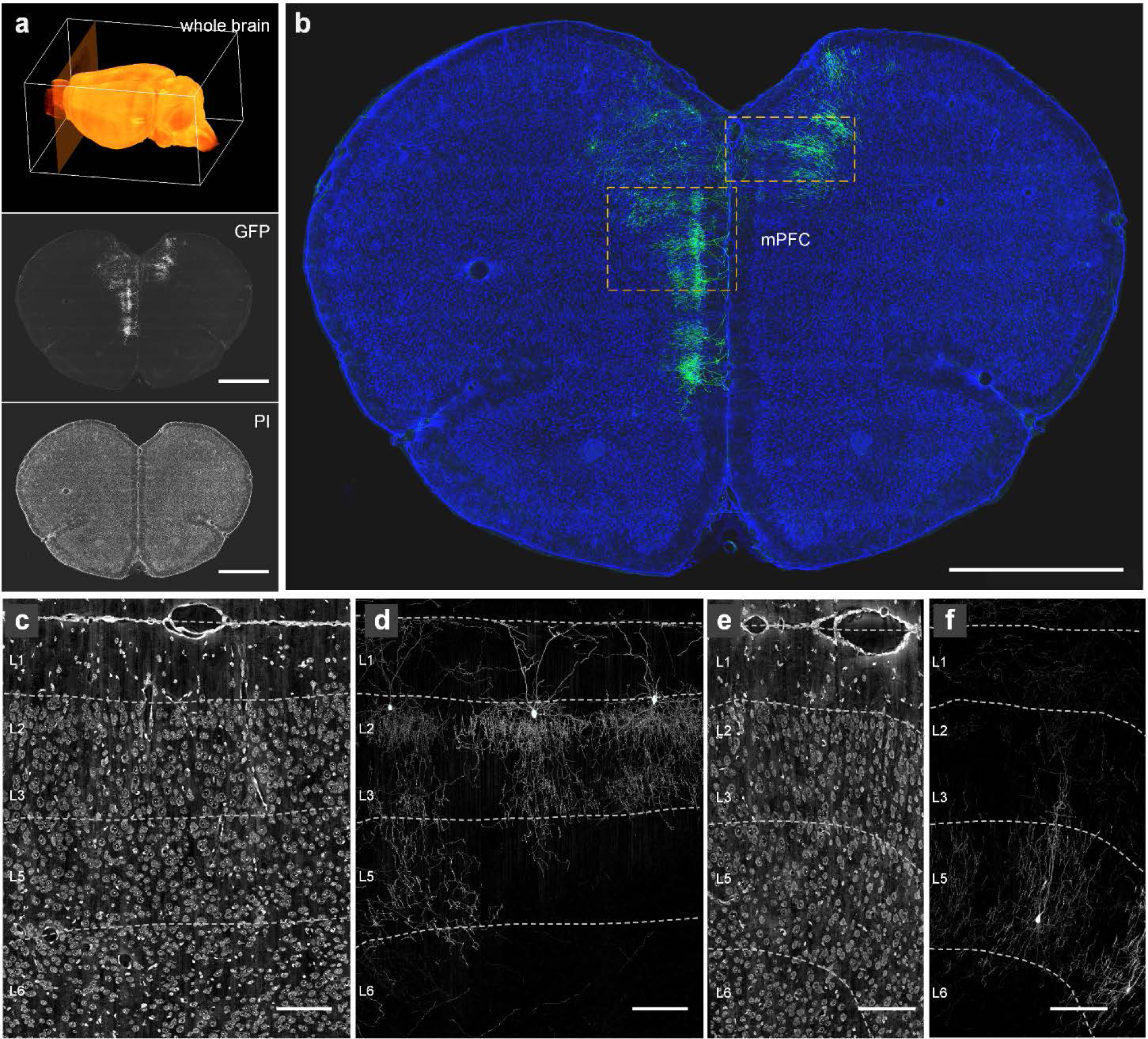
Areal and laminar distribution ofAACs revealed from whole-brain fMOST dataset. (**a**) A schematic of whole-brain coronal dataset collection (top) with an example of GFP channel (center) and PI channel (bottom) images. Scale bars: 1000 μm. (**b**) An example of the distribution of sparsely labeled AACs in mPFC.. Green: AAC morphology, 100 μm max-intensity projection. Blue: cytoarchitecture revealed by PI, 5 μm max-intensity projection. Scale bar: 1000 μm. (**c-d**) Laminar distribution of L2 AACs in mPFC. Enlargement of PI channel (**c**) and GFP channel (**d**) images from the left box in (**b**). (**e-f**) Laminar distribution of L5 AACs in mPFC. Enlargement of PI channel (**e**) and GFP channel (**f**) images from the right box in (**b**). Dashed lines in (**c-f**) indicate the layer boundaries. Cortical layers were discriminated based on cell body distributions in PI channel according to the Allen Mouse Brain Reference Atlas *(http://www.brain-map.org)*. Scale bars in (**e-f**): 100 μm

Despite the vexing diversity of cortical GABAergic neurons and the enduring debate on their classification scheme (DeFelipe et al., 2013), AACs were recognized as a bona-fide type soon after their discovery, largely based on their unique morphology and specific innervation of PyNs at AIS (Somogyi et al., 1982). Although the precise physiological action of AACs remains be elucidated (Lu et al., 2017; Szabadics et al., 2006; Woodruff et al., 2010), the defining feature is their specialization in regulating the spike initiation of PyNs. However, multiple variants of AACs have been found in several cortical structures (e.g. the hippocampus, piriform cortex, neocortex) and in different cortical layers that manifest different morphological features (Lewis and Lund, 1990; Somogyi et al., 1982; Taniguchi et al., 2013). This raises the questions of whether the cardinal AAC type consist of multiple “subtypes”, how should subgroups be defined, and what is the appropriate granularity.

We have previously captured cortical AACs through genetic fate mapping of neural progenitors of the embryonic medial ganglionic eminence (MGE) using the *Nkx2.1-CreER* driver line (Taniguchi et al., 2013). Conversion of transient Nkx2.1-CreER expression in MGE progenitors to a constitutive Flpase activity in AACs enabled postnatal viral targeting (He et al., 2016). By controlling CreER efficiency (i.e. tamoxifen dose), AAV injection volume and location, we were able to achieve specific, sparse, and complete labeling of AACs in defined cortical areas (**Fig. 2; Supplemental Fig. 1**). Here we analyzed AACs in the medial prefrontal (mPFC), primary motor (MC), and whisker-barrel somatosensory (SSC) cortex. We use the original nomenclature *axo-axonic cells (AACs)* (Somogyi et al., 1982) to refer to all GABAergic interneurons that innervate PyNs at AIS. Under this category, we use the term *chandelier cells (ChCs)* to refer to the subsets of AACs in the cerebral cortex (especially those in supragranular layers), whose axon arbors resemble the candle sticks of a chandelier light.

Following tissue resin-embedding and processing (Gong et al., 2016; Xiong et al., 2014), we used a dual-color fluorescence micro-optical sectioning tomography (dfMOST) system to image the whole brain samples at submicron resolution (**Fig. 1c; Movie 1**). The dual-channel capturing of neural morphology labeled by GFP and brain cytoarchitecture stained by propidium iodide (PI, red) were achieved by using a wide-field upright epi-fluorescence microscopy with a blue laser (488 nm) for fluorescence excitation and two separate TDI-CCD cameras for signal detection. Importantly, the PI channel provided each brain dataset with a self-registered Nissl like reference atlas of cell body distribution information, which allowed reliable delineation of cortical areas and layer boundaries (**Supplemental Fig. 1, 2**). Furthermore, the image contrast in PI channel was sufficient for the reconstruction of pyramidal neuron main dendrites which were used for identifying local laminar and vertical coordinates, readjusting cell orientation, and establishing a standardized platform for comparative analysis between cells in different cortical areas (**Supplemental Fig. 3)**.

From 11 whole brain dfMOST datasets, we completely reconstructed 62 AACs from mPFC, MC and SSC (**Fig. 3a; Supplemental Fig.14; Supplemental Table 1**). As axon arbors of AACs were extremely dense and complex, all AACs were manually reconstructed. This dataset represents the first set of complete and comprehensive AAC reconstructions in a brain region since their discovery 4 decades ago. The average length of AAC axons was ~2.1cm, average number of axon branches was ~1369, and average axon branch order was ~31. A major goal of our analysis is to define and discover AAC subtypes based on morphological features that inform connectivity, taking full advantage of the obligatory synaptic relationship between AAC axon terminals and PyN AIS. Our strategy was to examine the location and distribution of AAC cell bodies, their dendrite and axon arbor distribution, and their axon arbor geometry in the well-established coordinates of cortical laminar organization based on AAC postsynaptic targets – the PyNs (**Fig. 3b**).

**Figure 3.**
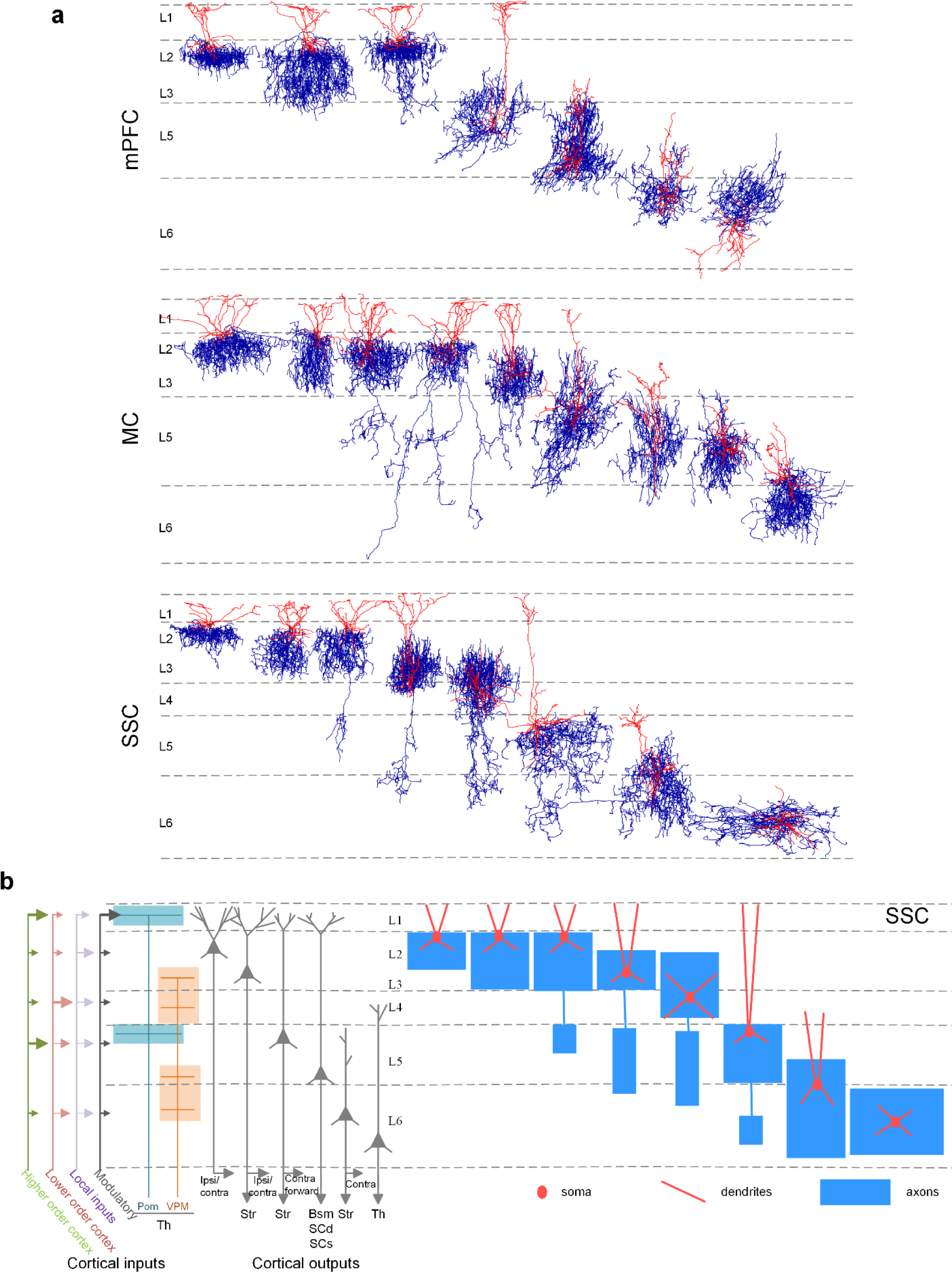
(**a**) Representative AAC single cell reconstructions in mPFC, MC and SSC. Cortical layers in each area were indicated by dashed lines. Black: soma body. Red: dendrites. Blue: axons. The orientation of each reconstruction was re-adjusted according to the local cortical vertical axis (see more details in **Methods** and **Supplementary Figure 3**). (**b**) Left: Scheme of the laminar arrangement of the input and output streams of SSC, in part rooted in the laminar organization of pyramidal neuron types with distinct projection targets. Right: a schematic of representative AACs in the SSC with characteristic laminar dendritic and axonal distribution patterns.

### AACs tend to localize at the borders between cortical layers

We have previously found that AACs are distributed across most if not all cortical areas and in multiple cortical layers (Taniguchi et al., 2013), but more precise description of AAC distribution has not been reported. The cellular resolution spatial coordinate information in the dfMOST datasets allowed unambiguous and quantitative localization of AACs. Within all three areas, the largest proportion of our reconstructed_AACs was located within the supragranular layers, with a major fraction at the layer 2 to layer 1 (L2/1) border (55%) and a much smaller set in layer 3 (5%) (**Fig. 2, 3; supplementary Fig. 1**). A significant portion of AACs were found in infragranular layers, both in layer 5 (L5 22%) and L6 (16%). We found one AAC in layer 4 of SSC. Interestingly, in most case, AAC somata tended to localize at the border between cortical layers, with prominent apical dendrites and basal axons (Figure 2, 3). At the areal border (defined by PI signal and cell distribution pattern), AAC axons appeared to restrict to one area and did not cross areas (**supplementary Fig. 5)**.

### AACs elaborate laminar restricted and predominantly apical dendrites that protrude dendritic spines

Almost all the reconstructed AACs elaborated prominent apical dendrites, while their basal dendrites were often sparse and restricted to the close vicinity of cell bodies (**Fig. 4**). The average span of apical dendrites of L2 AACs was 85.0 μm (90% of dendrite arbors horizontally cover 85.0 ±23.0 μm radial distance, mean ±SD, n=61) from soma (**Supplemental Fig.6**). In most cases, the apical dendrite extended within the one layer above the soma location (e.g. L1 for L2 AACs and L5 for L6 AACs). In several cases, L3 and L5 AACs extended apical dendrites all the way to the pia (**Fig. 3a; Movie 5 and Movie 6**). In particular, all L2 and some L3, L5 AAC dendrites appeared to tightly attach to the pia with thickened apical tufts; this is in contrast to many pyramidal neuron apical dendrites in L1 that do not reach near or adhere to pia surface (**Fig. 4a, Supplemental Fig. 4**). Interestingly, the apical but not basal dendrites of L2 ChCs sprouted filopodia-like slender dendritic spines, which were especially enriched in the upper half (68% in upper L1, the rest near L1/2 border) of L1 (**Fig. 4c-k**). The polarized dendritic arborization suggests that AACs receive most of their inputs from above their cell bodies. In particular, pia-attached AAC dendrites may recruit the most superficial L1 inputs and select or modify these inputs through dendritic spines.

**Figure 4.**
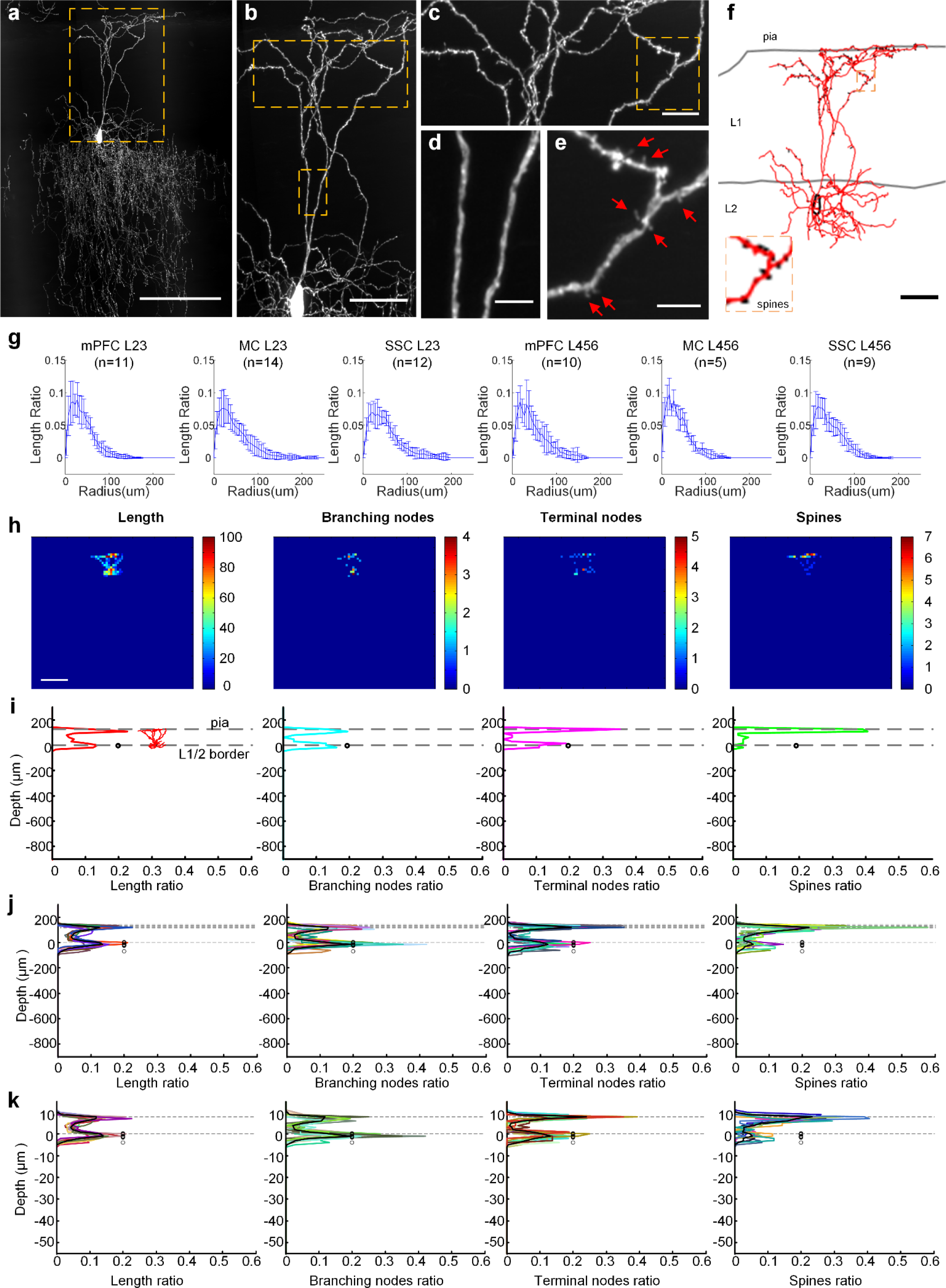
Characteristics of L2 AAC dendrites. (**a**) A representative L2 AAC in mPFC. 100 μm max intensity projection. Scale bar: 100 μm. (**b**) Dendrites of L2 AAC. Image was enlarged from the box in (**a**). Scale bar: 50 μm. Apical (**c**) and main dendrites (**d**) were enlarged from boxes in (**b**). Scale bars: 30pm (**c**) and 5 μm (**d**). (**e**) Spines (arrows) on the apical dendrites were enlarged from the box in (**c**). Scale bar: 5 μm. (**f)** Complete reconstruction of dendrites (red) and spines (black). Insert: enlarged from the box. Black circle: cell body. Gray lines: pia and L1/2 border. Scale bar: 50 μm. (**g**) Horizontal dendritic arbor distributions of up-layer (L2 and L3) and deep-layer (L4, L5 and L6) AACs in mPFC, MC and SSC. Data are mean ± SD. (h) An example of heat-maps showing the density distribution patterns of a L2 AAC dendritic arbor length (left), branching nodes (middle left), terminal nodes (middle right) and spines (right). Scale bar: 200 μm. (**i**) Single-cell density plots of a L2 AAC dendrites (same as in g) along the cortical depth. (**j**) Density plots of dendrites from 11 L2 AACs in mPFC. Different colors indicate different cells. (**k**) Normalized density plots of (**j**) based on pia and L1/2 border positions (see methods). Black circles in (**i-k)** indicate AAC soma positions in the coordinate. Dashed lines correspond to the place of pia (top) and L1/2 border (bottom). Dark black curve in (**j**) and (**k)** are averages of all the cells. Density value was presented by ratio.

### AACs elaborate laminar-stratified axon arbors, some with translaminar arbors

Although the characteristic shape and exquisite specificity of AAC axons have been recognized decades ago, few or none have been reconstructed to their entirety. We found that AACs axons arborized very extensively near the cell soma (below the soma for L2 AACs and both above and below the soma for other cortical AACs; **Fig. 5a-c; Supplement Fig. 7**). The average span of AAC axon arbors was 129.2 μm (90% of axon arbors horizontally cover 129.2±27.5 μm radial distance; mean± SD, n=61). In addition to the highly predominant local arbor (i.e. intralaminar), a significant fraction of L2 and L3 AAC axons (~74% of our L2 AAC reconstructions) further extended to the deeper layers (i.e. cross- and trans- laminar, **Fig. 3, 5b, d, Supplement Fig. 9; Movie 2, 3, 4**). In particular, translaminar axons of L2 AACs descended through intervening layers (e.g. L3-L5A in MC or L4 in SSC) before elaborating terminal branches with presynaptic boutons (**Fig. 5d**). This result suggests that, in addition to exerting powerful control over local PyN populations, some L2/3 AACs likely coordinate firing between local PyNs and a distant ensemble in an infragranular layer. Interestingly, we observed one L6 AAC with an inverted polarity – its dendrite extended below toward the white matter whereas the axon extended above toward L5 (**Fig. 3a, Supplement Fig. 13)**.

**Figure 5.**
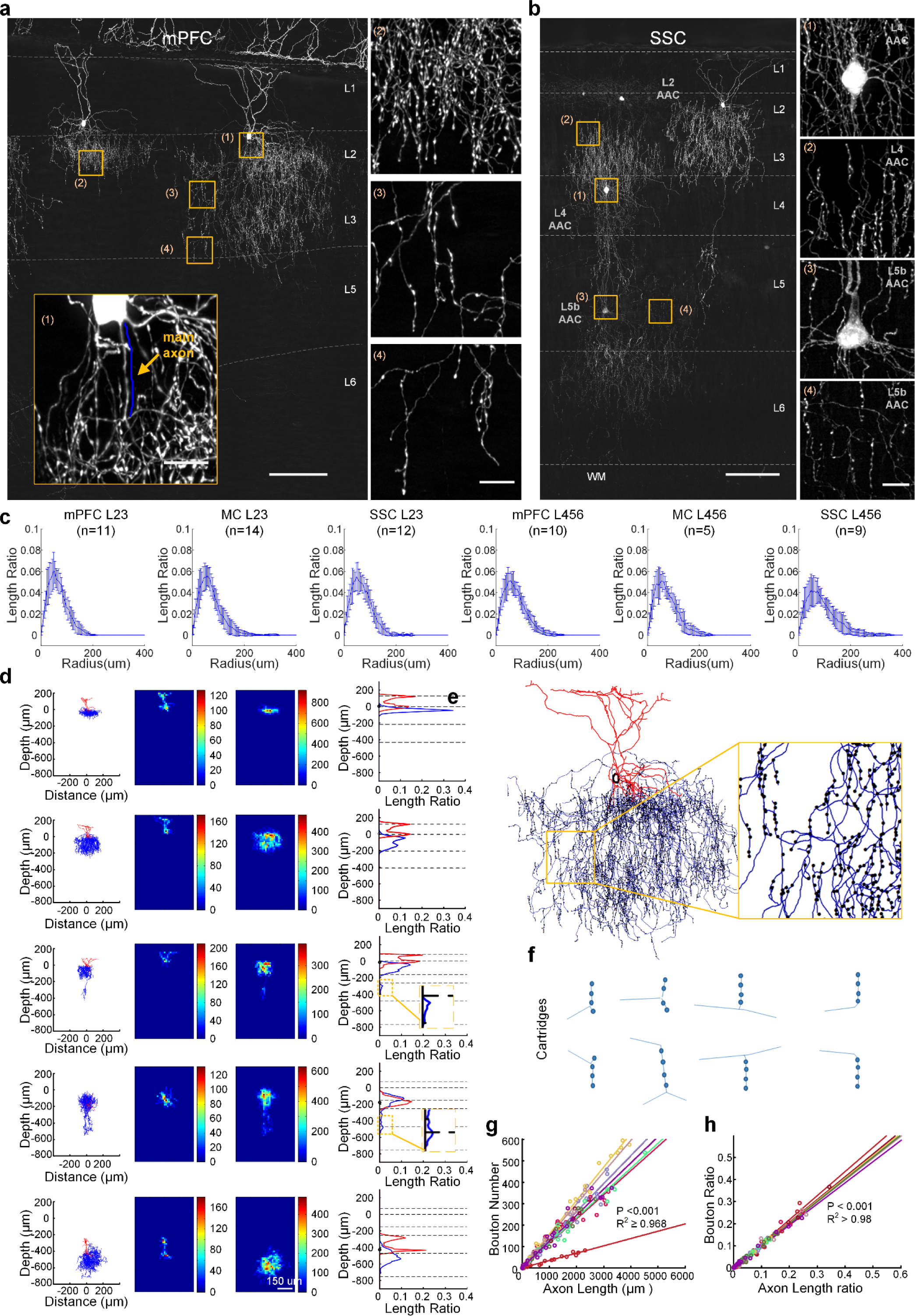
Morphology and distribution patterns of AAC axons in the neocortex. (**a**) A representative image (100 μm thick) of 2 nearby L2 AACs in mPFC. Inserts are enlarged images (boxes) showing the main axon (1; arrow) and typical axon cartridge clusters and individuals (2, 3, 4). Scale bars: 100 μm (low-mag image) and 10 μm (inserts) (**b**) A representative image containing nearby L2, L4 and L5 AACs in SSC. Enlarged L4 and L5 AACs were from the boxes in the left panel. Scale bars: 100 μm (left) and 10 μm (right). Dashed lines in (**a**) and (**b**) indicate cortical layers. (**c**) S Horizontal dendritic arbor distributions of up-layer (L2 and L3) and deep-layer (L4, L5 and L6) AACs in mPFC, MC and SSC. Data are mean ± SD. (**d**) Length density analysis of axons and dendrites from the AACs shown in (**a**) and (**b**). Left: projection of reconstructions (dendrites in red, axons in blue). Middle: heatmap of length density distribution of dendrites (middle left) and axons (middle right). Right: Length density plots of AAC dendrites and axons along cortical depth (y-axis). Dashed lines indicate layer boundaries. Insets in row 3 and 4 highlight axon branches in deep layers. (**e**) An example of axon bouton reconstruction of L2 AAC in mPFC. Insert: magnified view of the boxed region. (**f**) Axon cartridges that innervate PyN AIS can point upward, downward, or split from the middle. (**g-h**) The numbers of synaptic boutons correlate with axon length quantified by absolute value (**g**) or ratio (**h**).

### AACs consist of multiple subtypes distinguished by dendrite-axon distributions that reflect input-output connectivity patterns

The substantial variations in the location and morphology of AACs raise questions of whether they consist of anatomical “subtypes” and how subtypes can be resolved with biologically relevant granularity. As morphology is a proxy to and serves the purpose of connectivity, we first adopted a connectivity-guided approach to morphology-based AAC subtyping. Our analysis was based on the premise that at a mesoscale, establishing a synaptic connection requires the physical overlap between a presynaptic axon and its postsynaptic element within a specific anatomic location, i.e. an “anatomic parcel”, that represents the input or output component of a neural network (Ascoli and Wheeler, 2016); this tight spatial correlation often extends to the matching of fine-scale geometric features (e.g. presynaptic climbing fibers and postsynaptic Purkinje cell dendrites in the cerebellum). This obligatory correlation between pre- and post-synaptic elements, when framed in the context of circuit connectivity, provides a biologically relevant coordinate for morphological analyses.

The mesoscale correlation between pre- and post-synaptic elements is particularly identifiable and compelling for AAC and PyNs. Within the laminar architecture of the cortex, different types of PyNs that project to distinct cortical and subcortical targets are organized, to the first approximation, into different layers, and different sources of cortical and subcortical inputs are routed through laminar streams (Harris and Shepherd, 2015) (**Fig. 3b**). Importantly, the obligatory relationship between AAC axon terminals and PyN AIS represents a rare case where AAC axon distribution *alone* indicates connectivity to specific types of postsynaptic targets. Together these provide an inherent spatial coordinate system to register AAC position and morphology in the larger framework of cortical input and output streams (**Fig. 3b**). As the laminar arrangement of AAC dendrites recruit different input streams and the laminar stratification of axons mediate their output to separate PyN ensembles, we designed a AAC clustering analysis that emphasized the laminar density distribution of AAC dendritic and axonal arbors (**Fig. 6**) We excluded L3, L4 AACs (**Supplemental Fig 14)** from this analysis as there were few such examples (less than 4) in our current dataset.

**Figure 6.**
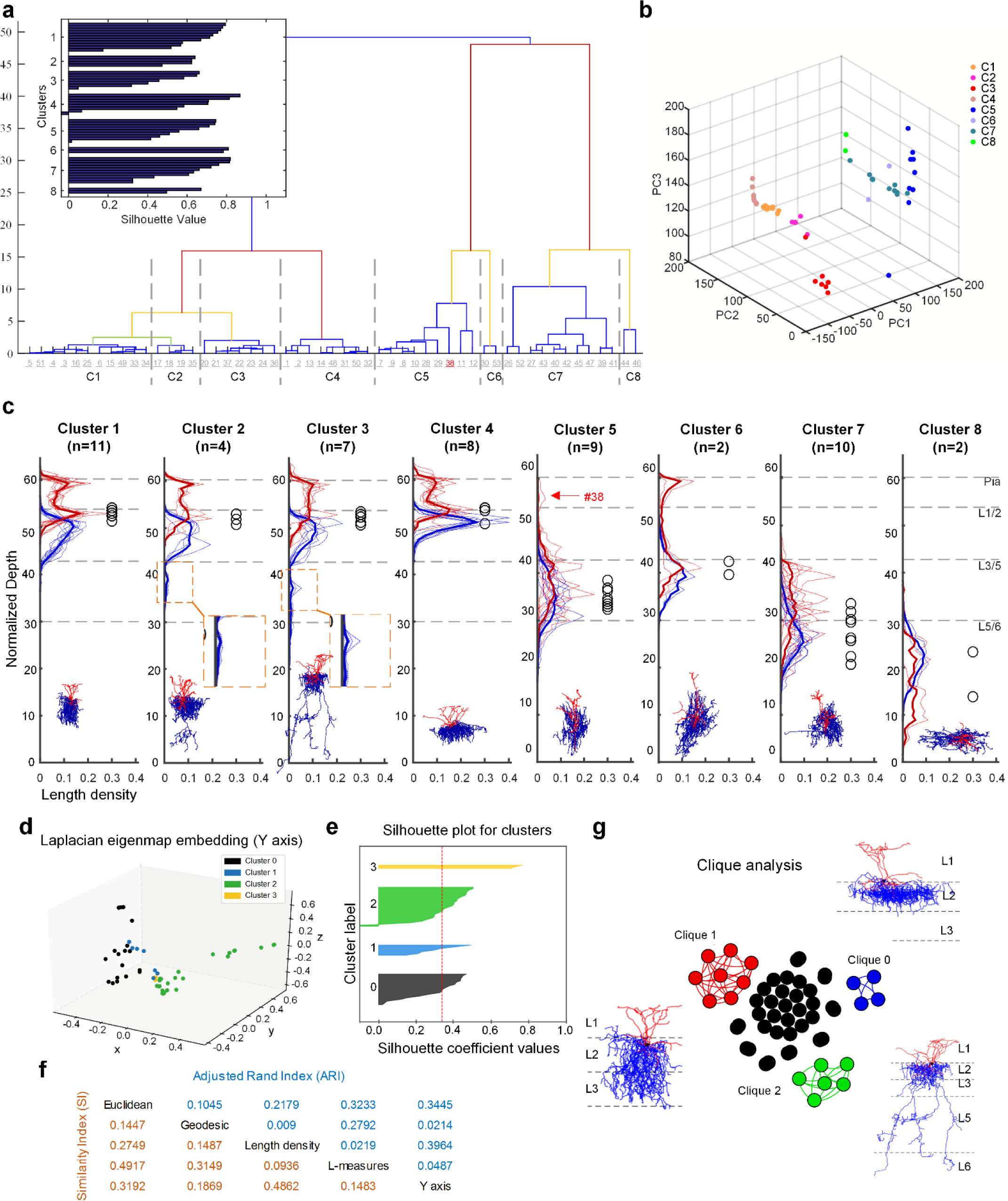
Hierarchical clustering of AACs based on cortical laminar density distribution of axons and dendrites. (**a**) Dendrogram of hierarchically clustered AACs (n=53). KL divergences (Kullback-Leibler divergence) of normalized arbor distribution functions along cortical depth were taken as the distance metric and furthest distance was taken as the linkage rule. See more details in Methods and Supplementary Figure? for the normalization procedures. Dashed lines correspond to the cutoff linkages of the identified 8 cell clusters. Insert: silhouette analysis of the 8 AAC clusters. (**b**) 3Dscattering plots of the 8 AAC clusters from (**a**) based on three principal components. (**c**) Axon (blue) and dendrite (red) length density distribution profiles of the 8 AAC clusters. Dashed lines: cortical layer boundaries. Black circles: soma body positions. Bold lines: average of all the neurons in each cluster. Note that cell #38 in cluster 5 has apical dendrites (arrow) reading L1 – a defining feature of cluster 6, but its lack of L3 axon branches (as it is located in SSC with a prominent L4) likely assigned it to cluster 5. (**d-g**) Clique analysis for the identification of robust AAC clusters. Clique analysis was conducted based on hierarchical clustering with 5 different metrics on AAC axons: Three persistent-homology based metrics, using 3 different ways of measuring distance from the soma, as scalar descriptor functions defined on the neuronal processes: euclidean, geodesic, and depth from cortical surface (“y-axis”), and the length density and L-measure metrics(Scorcioni et al., 2008) defined in the text (**d-e**). Laplacian eigenmap embedding of hierarchical clustering for the ‘y-axis’based metric (**d**) and other descriptors (**Suppl Fig. 16**). The selection of ‘K’ was based on silhouette analysis. Silhouette plot for K = 4 with y-axis as the input metric, thickness denotes sizes of clusters, red dotted line denotes average silhouette score, larger score means better clustering (**Figure 6e** and **Suppl Fig. 15**). The relations between the 5 metrics were quantified by Similar Index (SI) and Adjusted Rand Index (**f**). Three robust AAC clusters were identified based on the clique analysis (**Figure 6g** and **Suppl Fig.17**).

Based on brain cytoarchitecture information of dfMOST datasets, we normalized AAC dendrite and axon density distribution to a standardized cortex template **(Supplemental Fig. 11)**. Hierarchical clustering based on cortical laminar density distribution of axons and dendrites revealed multiple AAC subtypes grouped according to the laminar distribution of their cell body position and dendritic and axonal arborization. The identified L2 ChC clusters correspond to intra- (cluster 4), cross- (cluster 1) and trans- (cluster 2 and cluster 3) ChC subtypes. The axon arbors of cluster 3 extend both the L5 and L6 branches, but more dominantly innervate L5 **(Supplementary Fig 12)**. Cluster 7 AACs resided at L5 and L6 border (i.e.L6a), their dendrites were restricted in L5 and L6a and their axons arborized mainly in L6. Cluster 8 consisted of L6 AACs with intralaminar dendrite and axon arbors. These AAC subtypes likely receive different inputs and control different subsets of PyNs, and thus are distinguished by their circuit connectivity patterns. As the dendritic and axonal arbors of AACs are either present or not present in specific cortical layers to recruit inputs or innervate targets, respectively, AAC subtypes are likely unitary groups rather than subsets of a continuum. This was particularly apparent for two broad groups of L5 AACs, one extended short, L5-restricted apical dendrite and the other extended long, layer 1-reaching apical dendrites (**Fig. 3, Supplemental Fig 10**). We noted that this clustering method was not perfect as it assigned cell #38 to cluster 5, even though cell #38 extended apical dendrite to layer 1, as those characteristic to cluster 6 (**Fig. 6, Supplemental Fig. 8, 10**). In addition to these 8 clusters (**Fig. 6**), we detected 3 Layer 3 AACs (2 in SC, 1 in MC) with translaminar axon arbors and apical dendrite reaching layer 1 (**Supplemental Fig. 9; Movie 5**), 1 Layer 4 AACs in SC (**Movie 7**), and 1 inverted layer 6 AAC in mPFC (**Supplemental Fig. 13** show the projection image of L4 AAC).

**Figure 7.**
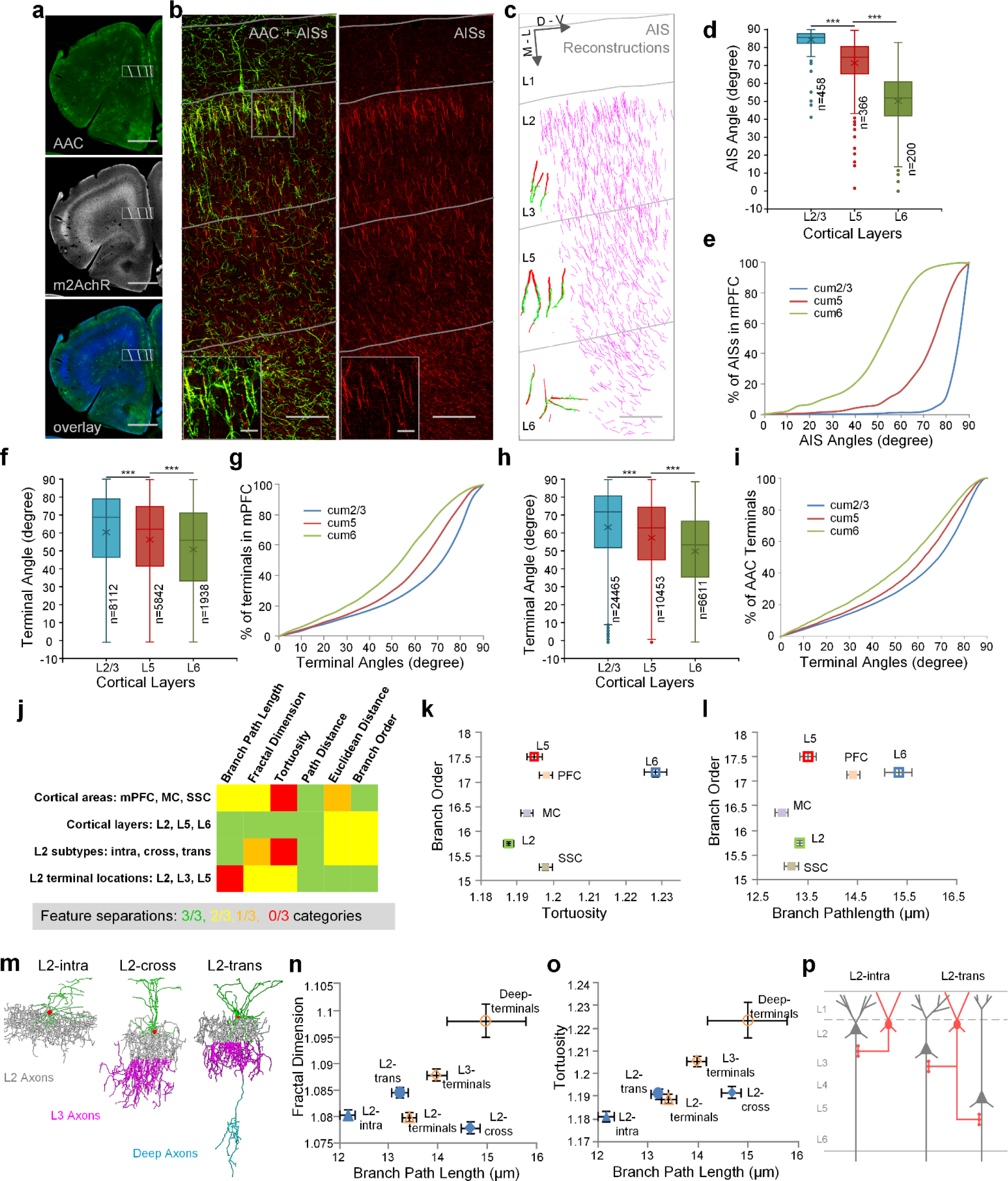
AAC subtypes revealed by axon terminal characteristics that correlate with AIS. (**a**) Distributions of AACs in mPFC (50 μm thick). AACs were labeled by the crossing of Nkx2.1-CreER mouse and Ai14 (LSL-tdTomato) mouse with low dose of TM induction at E18.5. Top: AACs (green) shown by the immunostaining of tdTomato. Center: cortical layers shown by the immunostaining of m2AchR. Bottom: color merged. Scale bar: 1000 μm. (**b**) AIS distributions in PrL (prelimbic cortex). Images were captured from the box in (**a**). Left: overlay image of AAC (green) and AIS (red). Right: immunostaining of AISs with Ankyrin-G. Inserts: enlarged images. Gray lines indicate layer boundaries. Scale bars: 100 μm (low-mag) and 20 μm (inserts). (**c**) AIS reconstructions (purple). Inserts: representative reconstructions of presynaptic AAC cartridge and postsynaptic PyN AIS pairs in L2/3, L5 and L6. Green: cartridges. Red: AISs. Scale bar: 100 μm. (**d-e**) Distribution differences of AIS angles among cortical layers in mPFC (***P < 0.0001, 95% confidence level, Kruskal-Wallis test followed by Dunn’s multiple comparisons test). Plots indicate median (full horizontal bar), mean (×), quartiles and range. AIS data (**d**) and cumulative plots (**e**) are from the reconstructions in (**c**). (**f-g**) The corresponding distribution differences of AAC axon terminal angles in mPFC (***P < 0.0001, 95% confidence level, Kruskal-Wallis test followed by Dunn’s multiple comparisons test). Plots indicate median (full horizontal bar), mean (x), quartiles and range. (**h-i**) Averaged distribution differences of AAC axon terminal angles across mPFC, MC and SSC (***P < 0.0001, 95% confidence level, Kruskal-Wallis test followed by Dunn’s multiple comparisons test). Plots indicate median (full horizontal bar), mean (x), quartiles and range. (**j**) Summary of axon terminal geometric features that separate AAC categories (cortical areas and cortical layers refer to somatic location). Green: statistically significant differences between all 3 pairs compared. Yellow: statistically significant differences between 2 of 3 pairs compared. Orange: statistically significant differences between 1 of 3 pairs compared. Red: no statistically significant differences. (**k-l**) Areal and laminar categories of AACs separated by axon terminal geometric parameters. Data are mean ± SEM. (**m**) Reconstructions of representative L2 AAC subtypes (L2-intra, L2-cross, L2-trans). (**n-o**) Axon terminal geometric parameters separate L2 AAC subtypes. Data are mean ± SEM.

Hierarchical clustering based on the laminar distribution of axon density alone has a potential shortcoming: it may over cluster or miss cluster two identical density profiles appearing at different layer depths. Further, low dimensional projections do not always show well-separated clusters and may need other indirect evidence about clustering in the high dimensional space such as silhouette plots. We therefore carried out a robust comparative analysis of morphological types using additional geometrical and topological characteristics of the neurons. For analyzing topological characteristics we used a recently developed framework employing persistent homology (Li et al., 2017) to derive a metric in the space of neuronal shapes. Briefly, this framework employs a descriptor function defined on the neurons, and a topological summary independent of neuronal location and orientation is derived from the descriptor function. We utilized three descriptor functions based on three different ways of measuring distances from the soma (Euclidean, Geodesic and Cortical Depth). In addition, we also used a community-standard metric(Scorcioni et al., 2008), employed on neuromorpho.org.

We performed hierarchical clustering employing each of these metrics, varying the number of clusters. By examining the overlap between the resulting clusters (ARI and SI indices, **Figure 6d, e**) we concluded that the metrics carry independent information about neuronal shape. We hypothesized that if a pair of neurons appears in the same cluster across all the metrics, this provides robust evidence that those neurons belong to the same morphological type. We thus proceeded by defining a graph in which each neuron is a node, and two nodes are connected if and only if they appear in the same cluster across all five metrics considered. This procedure produced a set of disconnected cliques (fully connected clusters). The three largest cliques corresponded to three robustly identified AAC cell types that are also visible in the hierarchical clustering using only the laminar density of the axons: the intra-, cross-, and trans- layer 2 AAC subtypes (**Figure 6g; Supplementary Figure 17**). A significant number of AACs could not be grouped into cliques, likely due to less than enough sample size. We hypothesize that with larger data sets, we will obtain similar robust cliques corresponding to other AAC subtypes for which preliminary evidence is provided by the hierarchical clustering shown in **Figure 6d-g**.

### AAC subtypes can be revealed by axon terminal geometry that correlates with that of postsynaptic AIS

In addition to the laminar stratification of axon arbors, AAC axon terminals in different cortical layers manifested different geometric characteristics such as orientation, tortuosity, path distance, and branch order (**Fig. 5e-h, Fig. 7**). As strings of AAC presynaptic terminals (i.e. “cartridges”) mostly align with the AIS of postsynaptic PyNs, we hypothesized that certain geometric features of AAC terminals reflect and correlate with those of the AIS. For example, the orientations of AIS in supragranular layers of mPFC were largely vertically aligned, but deviated substantially from this columnar orientation in infragranular layers (especially in L6; **Fig. 7a-c, Supplemental Fig. 12**). Consistent with this postsynaptic feature, AAC axon terminals in supragranular layers were also organized in predominantly vertical and parallel orientations, each largely straight and decorated with strings of presynaptic boutons (e.g. cartridges), which together earned them the name “chandelier cell”. In infragranular layers, on the other hand, the orientation of AAC terminals varied significantly with increased tortuosity that correlated with local PyN AISs (**Fig. 7d-i**). Interestingly, analysis of several geometrical features of AAC terminals properly grouped AACs according to areas, laminar positions, and L2 subtypes (**Fig. 7j**). In particular, several pairwise combinations of features classified AACs according to their areal, laminar locations and even the three subtypes within layer 2 (**Fig. 7k-o**). It is notable that AAC subtypes identified by axon local geometry are consistent with those identified by analyzing the laminar distribution of dendritic and axonal arbors (**Fig. 6**), both rooted in their connectivity to PyNs. Together, these results suggest that a connectivity-based framework of morphological analysis is informative in resolving the granularity of AAC subtypes.

## Discussion

As individual neurons are the basic building blocks of the nervous system, single neuron analysis is essential to reveal the true degree of cell diversity and principles of circuit organization. Single unit recording was key to discovering functional columnar organization in the cerebral cortex (Hubel and Wiesel, 1962), and single cell transcriptomics has facilitated understanding the molecular genetic basis of neuronal identity (Paul et al., 2017) and diversity (Shekhar et al., 2016; Tasic et al., 2016). As morphology is an intuitive depiction of neuron types that reflects their input-output connectivity, the visualization and quantification of complete single neuron shapes are necessary to identify and classify neuron types and deduce their anatomic relationships. However, the vast diversity, large spatial span, and vexing variations of mammalian neurons present unique challenges in cell labeling, imaging, and analysis. Recent advances in light microscopy begin to overcome the technical hurdle of submicron resolution imaging of the entire mouse brain using either wide field structured light illumination microscopy (i.e. MOST and fMOST; (Gong et al., 2013; Li et al., 2010) or fast scanning two-photon microscopy (Economo et al., 2016). In particular, the dual-channel dfMOST approach allows fast and simultaneous acquisition of both neural structures and their whole brain spatial reference at cellular resolution (Gong et al., 2016).

Another key requirement for reconstructing complete single neurons using light microscopy is sparse and robust labeling; and systematic labeling across neuronal populations is necessary to achieve comprehensive discovery of neuron types toward a cell census. Conventional transgenic approach lacks specificity and sparseness. Although viral vectors can achieve sparse labeling of distal axons (Economo et al., 2016), their limitations include 1) dense labeling of local collaterals that are difficult to reconstruct, 2) lack of specificity to local interneurons, and 3) lack of orthogonal information (e.g. molecular markers) to further restrict labeling and help interpret morphological variations in cell type identification. Our combinatorial genetic strategy overcomes these limitations. We engage multiple cell features to target subpopulations defined by gene combinations, lineage, birth time and anatomy (He et al., 2016). We further incorporate inducible and viral methods to achieve reliable single cell labeling (**Fig. 1b**), which enables “saturation screening” of morphological types or subtypes within the subpopulation. Although here we have not reached saturation screening of cortical AACs, as L4 and inverted L6 AACs were detected only once in our dataset, the approach demonstrates unprecedented specificity and comprehensiveness to one of most rare cortical cell types known to date. Iterations of this labeling scheme through systematic generation of mouse driver lines (Harris et al., 2014) will enable comprehensive targeting of neuron types as has been demonstrated in genetic targeting in *Drosophila* (Aso et al., 2014; Jenett et al., 2012) Thus together with scRNAseq, gSNA provide an orthogonal quantitative and scalable single neuron analysis platform. Currently, the bottleneck of the gSNA is single neuron reconstruction, which mostly relies on manual procedures. Future innovation in machine learning-based automatic reconstruction algorithms may increasingly overcome this limitation (Peng et al., 2015; Peng et al., 2017).

The goals of single neuron anatomy are to identify and catalog cell types and, ultimately, to inform the study of cell function through inferring connectivity. With the prospect of increasing throughput in single neuron reconstruction, a pressing issue is how to extract biologically relevant information from morphology datasets. Traditional approaches deploy a large set of geometric and topological metrics (e.g. (Petilla Interneuron Nomenclature et al., 2008)) to quantify single neuron morphology in isolation often without a proper spatial coordinate and circuit context; these analyses are mostly ineffective in parsing neurons into reliable and biologically informative groups. As morphology is a proxy to and serves the purpose of connectivity (Seung and Sumbul, 2014; Sumbul et al., 2014), we have adopted a connectivity guided approach to morphological analysis. This approach is based on the premise that, although single neuron shape by itself does not contain information about presynaptic sources and postsynaptic targets, such information can be extracted, to a varying degree, if neuron morphology can be registered and analyzed in an appropriate spatial coordinate that reflect local and/or global connectivity patterns. Indeed, the inherent polarity of dendrites and axons ensures that their distribution and geometry reflect the input source and output targets in the corresponding spatial domains, or “anatomic parcels” (Ascoli and Wheeler, 2016). Although the precise identity of input and output elements cannot be inferred from spatial location alone, anatomic parcels based on decades of classic studies provide significant information to include and exclude pre- and post-synaptic elements and thus to infer possible as well as impossible connectivity. This analysis framework is likely to recognize seemingly “subtle” morphological variations (e.g. translaminar dendrite or axon branches of AACs) which yet have significant impact on connectivity and thus cell function. In this context, the dfMOST datasets, which allow automatic registration of single neuron morphology into proper global and local coordinates at cellular resolution within the same brain (Gong et al., 2016), is key in analysis strategies to identify and distinguish cell types and to inform connectivity. In analyzing AAC morphology, for example, the precise cell distribution information of the dfMOST dataset is crucial to derive and normalize laminar coordinates in different cortical areas, which enabled areal and laminar comparisons and inferences of input-output connectivity patterns that distinguished AAC subtypes. Our results indicate that high resolution morphology dataset *alone*, when registered within proper spatial coordinates that reflect brain circuit organization, contain rich anatomical information on cell identity and connectivity. This analysis framework should apply to projection neurons as dfMOST datasets contain brain wide information on anatomical parcels that will inform the potential synaptic targets of long range axon branches. Therefore, light microscopy-based high throughput single neuron anatomy will likely provide substantial information and insight on cell type diversity and mesoscale connectivity in the mammalian brain.

A recent study suggests that cardinal GABAergic neuron types are distinguished by their input-output synaptic communication patterns encoded in transcription profiles (Paul et al., 2017). This synaptic communication framework of neuronal identity may integrate anatomical, physiological, molecular, and functional descriptions of neuron types (Lerner et al., 2016; Paul et al., 2017). Beyond cardinal types, finer division into subtypes may better represent and explain the intricacies of neural circuit organization (Zeng and Sanes, 2017), but there is no consensus and mechanistic basis on the granularity and boundary of neuronal subtypes. Our results on cortical AACs suggest that input-output connectivity, reflected in cell morphology, is likely a key arbiter of neuronal subtypes. Differential gene expression in supragranular or infragranular AACs in the frontal cortex (Paul et al., 2017) is consistent with this interpretation. We therefore suggest that synaptic communication patterns may distinguish neuronal subtypes as well as cardinal types. It is notable that AAC subtypes, when registered along cortical laminar coordinates, appear to manifest a degree of stereotypy and fine granularity that is similar to those of retinal bipolar cell subtypes registered upon the much finer coordinates of retinal sub-laminae (Shekhar et al., 2016). A true saturation anatomical analysis of cortical AACs will most likely reveal additional subtypes. While the division of retinal bipolar subtypes is further supported by molecular, physiological and functional evidence (Euler et al., 2014), the division of AACs subtypes based on anatomy needs to be substantiated by orthogonal datasets, including their physiological connectivity (Lu et al., 2017), gene expression profiles (Paul et al., 2017), and possibly developmental genetic basis (Taniguchi et al., 2013).

## Author contributions

Z.J.H. and Q.L. conceived and organized the study. H.G. supervised and organized the imaging, reconstruction, and analysis. X.W. performed viral labeling and resin-embedding experiments. J.T. developed genetic viral targeting of AACs. F.Y., T. Y., X.W. contributed to the dfMOST system design, maintenance and data capturing under the supervision of S. Z. B.S.W. contributed to AIS labeling and confocal imaging experiments. Y. J., Y. L., X. W. performed the image preprocessing and format transformation. S.J., X.J., X.W. conducted neuronal reconstructions and delineation of cortical layer boundaries. S.J., X.W., Z.X. performed data analysis and modified the methods. X.W., Z.J.H., H.G., P.M., M.A.A., G.A. A., Y.W., D.W. analyzed the data. Z.J.H., X.W., P.M. wrote the manuscript with comments from all authors.

## Acknowledgement

This work was supported by the National Natural Science Foundation of China (Nos. 61421064, 91432105) and Director Fund of WNLO (Q.L. H.G, S.Z.), 5R01MH094705-04 and CSHL Robertson Neuroscience Fund (Z.J.H.), R01NS39600 and Burroughs Wellcome Fund Collaborative Research Travel Grant (G.A.A.), 1R01 EB022899-01 (Y.W., P.M.), and a Crick-Clay Professorship (P.M.). J.T. and B.S.W were supported by NRSA pre- and post-doctoral fellowships, respectively.

## Tables

**Table 1**. Complete cell list and Neurolucida dataset on morphometry

## Movies

**Movie 1**. dfMOST imaging of viral labeled AACs at single-axon resolution. 100 μm max intensity projection of original GFP channel images without any imaging processing.

**Movie 2**. Nearby L2-intra and L2-cross AACs in mPFC.

**Movie 3**. Reconstructions of nearby L2-intra and L2-cross AACs in mPFC.

**Movie 4**. Reconstructions of nearby L2-intra and L2-cross AACs in SSC.

**Movie 5**. Reconstructions of nearby L3 and L5-intra AACs in MC.

**Movie 6**. A L5a (L5-cross) AAC in mPFC.

**Movie 7**. Nearby L4 and L6a (L5/L6 border) AAC in SSC.

## Materials and Methods

### 1. Experimental animals and low dose TM induction

To achieve sparse and specific targeting of AACs across neocortical areas, we crossed *Nkx2.1-CreER* mice (The Jackson Laboratory stock 014552) with *Rosa26-loxp-stop-loxp-flpo (LSL-Flp)* mice (The Jackson Laboratory stock 028584). At postnatal day 0 (P0) or day 1 (P1), we intraperitoneally induced each pup with low dose of tamoxifen (TM, 0.25 mg per pup). Tamoxifen stock solution (5mg/ml in corn oil) were prepared beforehand. Sparsely targeted AACs would express FlpO constitutively (He et al., 2016).

For immunostaining experiments, we crossed *Nkx2.1-CreER* mice with *Rosa26-lox-stop-lox-tdTomato (Ai14)* mice (The Jackson Laboratory stock 007905). To ensure embryonic day 18.5 (E18.5) TM inductions, Swiss Webster or C57B6 females (Taconic) were housed with Nkx2.1CreER;Ai14 (ht/homo) males overnight and females were checked for vaginal plug by 9am the following morning. At E18.5, pregnant females were given oral gavage administration of TM (dose 3mg / 30g of body weight) for sparse labeling of AACs. AACs are labeled with tdTomato. Genetic hybrids of C57B6 and Swiss Webster animals were used in these experiments. All animal breeding and surgical experiments were approved by the Institutional Animals Care and Use Committee (IACUC) of Cold Spring Harbor Laboratory or the Institutional Animal Ethics Committee of Huazhong University of Science and Technology.

### 2. Stereotaxic virus injection

Flp dependent AAV-fDIO-TVA-GFP (TVA: avian glycoprotein EnvA receptor) cassette was assembled and cloned using standard molecular cloning protocols with restriction enzymes from New England Biolabs. TVA-GFP *(pAAV-EF1a-FLEX-GT)* was a gift from Ed Callaway (Addgene plasmid # 26198). The cassette was subcloned into *AAV-Ef1a-FD-YFP-WPRE* (a gift from the Deisseroth laboratory, Stanford University) using NheI and AscI cloning sites(Fenno et al., 2014). All constructs were sequenced to ensure their fidelity and proper reversed orientation of the inserts, and packed into AAV8 viral vectors with titers ranging from 1.0 × 10^12^ to 2.4 × 10^12^ pfu from the UNC Vector Core *(Chapel Hill, North Carolina*).

For stereotaxic injection, post-weaned animals (3 to 4-week-old) were anesthetized by intraperitoneal injection with katamine and xylazine (ketamine:100 mg/kg, xylazine: 10 mg/kg in saline), and then were fixed in a stereotaxic headframe *(Kopf Instruments Model 940 series)* for the identification of the coordinates of mPFC, MC and SSC areas based on the Allen Mouse Brain Reference Atlas *(http://atlas.brain-map.org)*. Each animal received bilateral injection in mPFC, MC and SSC areas (6 injection sites per mouse). At each site, we injected 100 nl virus with Nanoliter 2010 Injector *(World Precision Instruments)*. And we let virus express more than 21 days for strong labeling. The membrane tagged labeling by TVA-GFP fusion significantly improved the labeling of fine axon terminal. The stereotaxic coordinates are: mPFC (A/P: 1.98 mm, M/L: ±0.5 mm; D/V: 1.5mm depth from pial surface), MC (A/P: 0.5 mm, M/L: ±1.5 mm; D/V: 0.5 mm) and SSC (A/P: -1.5 mm, M/L: ±3.0 mm; D/V: 0.5mm).

### 3. Perfusion and whole-brain resin embedding(Gang et al., 2017).

Mice were deep anesthetized by overdose of ketamine and xylazine, and then intracardially perfused with 0.01M PBS (Sigma-Aldrich Inc., St Louis, MO, USA) and 4% paraformaldehyde (PFA, Sigma-Aldrich Inc., St Louis, MO, USA). After brain dissection and about 18 hours of post-fixing in 4% PFA, brain samples were rinsed in 0.01M PBS overnight. Then samples were dehydrated in graded series of ethanol (with distilled water): 50% ethanol (2h, 3 times), 75% ethanol (2h, 1 time), 100% ethanol (2h, 2 times). After dehydration, we replaced ethanol with graded series of xylene (with pure ethanol): 50% xylene (2h, 2 times) and 100% xylene (2h the first time, and then overnight). We then infiltrated samples in graded series of Lowicryl HM20 resin (in xylene): 50% HM20 (2h), 75% HM20 (2h), 100% HM20 (2h, 2 times), 100% HM20 (2 days). After resin infiltration, samples were heat-polymerized at 50 °C for 8 hours in a vacuum oven. All dehydration and infiltration procedures were treated at 4 °C. All solutions were prepared in weight.

Note: During wide-field based dfMOST imaging, autofluorescence produced by lipofuscin in the resin-embedded brain tissue often interfered with image contrast. Swiss Webster mice, especially after 2months of age, usually express more lipofuscin compared with C57/BL6 mice. To reduce the effect of lipofuscin, all animals in this study were sacrificed around P51-P54.

### 4. Whole-brain dual-color fMOST (dfMOST) imaging

Plastic embedded brain samples were mounted on a metal base and then installed under a dual-color fluorescence micro-optical sectioning tomography (dfMOST) system for whole-brain imaging. The dfMOST system is a wide-field block-face imaging system. A blue laser (488nm) was used as the excitation light source with two separate TDI-CCD cameras for signal detection. This system runs in a stripe scanning mode (X axis) and combines with an afterward image montage to realize the centimeter-scale coronal data acquisition(Yang et al., 2015). A precision motorized XYX stage is used to conduct imaging scanning, areal expansion and ultra-thin sectioning by a diamond knife (Li et al., 2010). The high throughput and high resolution imaging of fluorescent protein labeled samples is realized with chemical sectioning (Xiong et al. in preparation). Following each scanning of one coronal plane (X-Y axes), the sample was sectioned to remove the top layer (Z axis), and then imaged again. The imaging-sectioning cycles were performed automatically with 1.0 μm Z steps until whole brain was imaged. The resin-embedded GFP fluorescence were well preserved through chemical reactivation (Xiong et al., 2014) provided by adding Na2CO3 in the imaging water bath (0.05 M, PH = 11.4).

We used a 60X water immersion objective (NA 1.0) for imaging, which provided the system with submicron resolution at 0.2 × 0.2×1 μm voxel sampling rate for each whole-brain dataset. High resolution and high density sampling rate greatly facilitated our cell reconstruction procedures and are especially necessary for reconstructing dense neural arborizations and fine structures (such as axon boutons and spines).

The red channel was used to capture the whole brain cytoarchitecture which was counterstained by propidium iodide (PI)(Gong et al., 2016). PI dyes were dissolved in the imaging water bath, thus stained the exposed DNAs and RNAs on the tissue surface (also see **Supplementary Fig. 2**).. The staining occurred within thus was in real time. The 488 nm laser was strong enough for PI excitation. The counterstained cytoarchitecture provided a self-registered Nissl like brain atlas for the GFP channel and was used for the identification of cortical areas and layers. Furthermore, the image contrast in the PI channel was sufficient for identification and reconstruction of pyramidal neuron main dendrites (**Supplementary Fig. 2** and **Supplementary Fig. 3**). Weakly stained or unstained tubular cellular objectives, such as blood vessels and pyramidal main dendrites, can be seen in good contrast in PI channel.

### 5. Single cell reconstruction and layer boundaries discrimination.

To reconstruct sparsely labeled single ChCs from the whole-brain image datasets (~ 8 TB), we transformed TIFF format raw images series to LDA type(Li et al., 2017a). We then used Amira software *(v 5.2.2, FEI, Me’rignac Cedex, France)* to load the LDA data and identify cells for initial reconstruction. We only chose cells with highly characteristic axon terminal cartridges which were true ChCs (~30% GFP-labeled neurons were not the ChC type). The areal and laminar location of selection of cells were identified based on the cytoarchitecture provided by PI staining according to Allen Mouse Brain Reference Atlas. All initial reconstructed cells were saved in SWC format.

The arborization of a complete single ChC was extremely dense: the average length of AAC axons was ~ 2.1cm, the average number of AAC axon branches was ~ 1369, and the average axon branch order was ~ 31. Only manual procedure was feasible to reconstruct cells with such arbor complexity. Each AAC took up to one week to complete by one person.

To ensure all AAC reconstructions were correct and complete, we carried out revisions on each initial reconstruction in Neurolucida360 software *(Neurolucida, MBF Bioscience, Williston, VT)*. Since neurolucida 360 was not compatible with the reading of SWC format files and couldn’t hold TB-size image datasets, we transformed all the SWC files to Neurolucida ASC format using the Neuronland software (Neuromorpho.org), and we cropped smaller image stacks (GB-size) of GFP and PI channels from the whole brain data sets. The Cropping areas were based on the coordinates calculated from the initial SWC reconstructions.

Cortical layer boundaries were reconstructed in the co-registered PI channel in Neurolucida 360. Laminar positions were discriminated based on cell body distributions according to the online version of Allen Mouse Brain Atlas *(http://www.brain-map.org)*. 5μm max intensity projections of PI images were used to better show the cell body distributions (**Supplementary Fig. 2**).

### 6. Adjusting the orientation AACs to the vertical axis of local cortical column

To identify the local vertical axis of cortical depth, we randomly reconstructed a few pyramidal main dendrites near the reconstructed AAC cell body in PI channel. We took the main direction of pyramidal neuron apical dendrites near the AAC cell body as the cortical column vertical axis. We first randomly reconstructed a few pyramidal apical dendrites. We then centered all the traced pyramidal dendrites and calculated their main orientation by Principal Component Analysis (PCA). Using this orientation as the proxy of cortical vertical axis, we re-orientated each AAC reconstruction using the MATLAB software. More details in **Supplementary Fig. 3**.

### 7. Length density analysis

<u>Length density analysis of AAC morphology</u>

Length density analysis of dendrites and axons were performed using custom Matlab routines(Yamawaki et al., 2014). Briefly, for each orientation-readjusted AAC, we set the soma center as origin of coordinate. The neuronal arbors were divided into 15 μm × 15 μm grid space in the xy plane, and the arbor length in each grid were summed covering the whole z direction. The distribution pattern in coronal plane (i.e. xy plane) was represented in heatmap. Length density profile along the cortical depth direction (i.e. y-axis) were plotted to quantify the laminar distribution pattern by integrating fiber length along x direction from heat-map. Similarly, length density profiles along x-axis (middle-lateral) and z-axis (anterior-posterior) were plotted to quantify the horizontal distribution patterns (**Supplementary Fig.7 b-f**). To make easy comparison, we normalized profile by dividing the fiber total length of the cell (length ratio). Layer boundaries were also plotted in the length density figures (dashed lines). Their positions in the y-axis was the average coordinates of all the contouring points covering the neuron arbor extent in x direction (**Supplementary Fig. 11**).

<u>Normalized Length density distribution on a standard cortex template</u>

For comparative analysis among AACs from different brain areas and samples, we normalized the laminar distribution of AAC axonal and dendritic arbors to a standard cortex template (y-axis only). In the neocortex, only SSC has L4 compared with mPFC and MC, and the L6-WM (white matter) border in mPFC is usually not discernable in the coronal plane. And the thickness of the same layer in different cortical areas and even subareas can be different. To address these issues, we performed normalization based on the thickness of each layer, rather than using the distance from pia to WM. The number of laminar arbitrary units (AUs) been used for subdividing each layer was decided based on the average thickness of each layer from all cells (L1: 100.04 μm, L2/3: 180.52 μm, L5: 215.33 μm, n=53). Here we kept the dividing size to be around 15 μm to match with the unnormalized length density analysis. Thus, the numbers of laminar AUs for dividing L1, L2/3 and L5 are 7 AUs, 12 AUs and 14 AUs respectively. According to these parameters, as shown in **Supplementary Fig. 11**, axon arbors were subdivided with different intervals for L1, L2/3 and L5. For the arbors above L1 and below L5 (L6), we used the dividing intervals of L1 and L5 respectively. Since the axons of most AACs did not innervate L4 (except one L4 AAC), we removed L4 length density data for all the AACs in SSC. Based on this method, we could get normalized length density distribution curves of dendrites and axons for each cell from all the three cortical areas.

### 8. Cluster analysis

Based on normalized distribution of neural arbor length density along the y-axis (cortical depth), 53 AACs were hierarchically clustered using a weighted KL divergence (KLD_w_, symmetrized(Don H. Johnson, 2001)) as the distance metric and the furthest distance as the linkage rule. A weight coefficient λ was defined as the ratio of average axon length to total axon and dendrite length across all neurons. The KLDw matrix was calculated by multiplying the axon distribution by the length based weight λ and the dendrite distribution by (1 – λ). That is KLD_w_ = λKLD_axon_ + (1 – λ)KLD_dendrite_. Based on the clustering dendrogram of KL divergence, 53 AACs can be grouped to different clusters. Corresponding silhouette analysis was done based on the cutoff linkages used in the clustering.

### 9. Clique analysis

To robustly classify the AACs we did a comparative clustering study across five different metrics, to find neuronal groups that clustered together irrespective of metric utilized.

Three of the metrics were derived from topological considerations described in (Li et al 2017)(Li et al., 2017b). This methodology starts by defining a “descriptor function”, which is a scalar valued function defined on the axons and dendrites of each neuron. The procedure then computes a topological signature known as the persistence diagram for each neuron based on the descriptor function. Finally, the distance between two neurons is defined by computing a suitable metric between the persistence diagrams. The persistence diagrams are by definition invariant to rigid translations and rotations, and may have further invariances. Three of the metrics were defined by using three different descriptor functions, in each case defined as a suitable distance from the soma to the point on the neuron. The three distance functions used were Euclidean distance, Geodesic distance along the neuron, and distance along the normal to the cortical sheet (we denote this the “y-axis” for brevity).

In addition, we used a metric defined by taking KL distance between the histograms created by projecting the neuronal processes onto the y-axis (“length density”), and finally a community-standard metric, the L-measure(Scorcioni et al., 2008), used on neuromorpho.org.

How related are these metrics? To answer this question, we performed hierarchical clustering using each of the metrics, fixing the total number of clusters to be K. In general the different metrics produced different sets of clusters. We compared the sets of clusters across two metrics, using the Adjusted Rand Index (ARI), and the Similarity Index(Bohland et al., 2009) (SI) defined in Bohland et al (http://journals.plos.org/plosone/article?id=10.1371/journal.pone.0007200) to compare different parcellations of brain atlases. In each case, these indices lie between 0 and 1, with 1 corresponding to perfect correspondence between two sets of clusters. We found (**Fig 6f**) that both indices were generally closer to 0 than to 1, indicating that these metrics measured independent geometrical/topological characteristics of the neurons. Thus, if neurons were grouped together by all five metrics, we would gain confidence that they were robustly classified into these clusters.

To perform this robust classification we used the following method: (i) first, we carried out hierarchical clustering using each of the metrics, with a fixed number K of clusters, (ii) We then defined an undirected graph G with each node corresponding to a neuron. The edge between two neurons is either 1 or 0 based on whether the neurons clustered together or not as described below. (iii) We then looked for disjoint cliques (in a clique, each node is connected to every other node in the clique; intuitively, a clique constitutes a set of very similar neurons). These disjoint cliques were our robust clusters.

Let the number of metrics be M (=5 in our case). We introduced a parameter N that controlled the edge weights of the graph G as follows: If two neurons belonged to at the same cluster for at least N of the M metrics, then we gave that edge a weight 1, otherwise we gave it a weight 0. Thus, the graph G was a function of two parameters K,N. We then looked for maximal cliques in G(K,N). For N<M, the maximal cliques in G were not in general disjoint, however for N=M the cliques can be shown to be disjoint. Consider the relation between two neurons given by an edge in G(K,M). This relation is transitive: if two pairs of neurons (N1,N2) and (N2,N3) are connected, then (N1,N2) must belong to the same cluster across all metrics, as well as neurons (N2,N3). It follows that (N1,N3) must also belong to the same cluster (of which N2 is a member). This transitivity guarantees the disjointedness of the maximal cliques: If two cliques share a vertex, then the two cliques must be identical. Thus, we considered only G(K,M) and found the disjoint maximal cliques. In our case M=5. We selected K by examining the average silhouette scores of the clusters versus K (http://scikit-learn.org/stable/auto_examples/cluster/plot_kmeans_silhouette_analysis.html#sphx-glr-auto-examples-cluster-plot-kmeans-silhouette-analysis-py). <u>Finally, performing clique analysis on G(K=4,N=M=5), we found 3 cliques with size greater than 2 (with sizes 4,6 and 8 respectively;</u> **Fig 6g**). <u>These cliques were the output of our robust clustering analysis, and exemplars from each clique are showin in</u> **Fig.6g**.

### 10. Immunostaining

Adult animals (P45-P60) were perfused with 4% PFA in PBS. The brains were removed and post-fixed overnight in the same fixative. Coronal brain slices were sectioned at 75 um thickness via vibratome. Sections were blocked with 10% normal goat serum in 0.5% Triton in PBS for an hour and then incubated overnight with primary antibodies diluted in blocking solution at room temperature. Primary antibodies used were: rabbit polyclonal RFP (1:1000, Rockland) for labeling AACs, mouse monoclonal Ankyrin-G ( 1:500, Neuromab) to label pyramidal axon initial segments (AIS), and rat monoclonal muscarinic Acetylcholine receptor m2 (m2AChR) (1:500, Millipore Sigma) to discriminate L3/5 and L5/6 boundaries in mPFC. Sections were subsequently washed and incubated with the appropriate fluorescently-conjugated secondary antibodies diluted in the same buffer for 3 hours at room temperature. Secondary antibodies used were: Alexa Fluor 488 goat anti-rat (1:500, Invitrogen), Alexa Fluor 594 goat anti-rabbit (1:500, Invitrogen), and Alexa Fluor 647 goat anti-mouse IgG2a (1:500, Invitrogen).

### 11. Code availability

All custom codes used in this study are available from the corresponding author upon reasonable request.

### 12. Data availability

The data that support the findings of this study are available from the corresponding author upon reasonable request.

